# Cardiac afferent activity modulates early neural signature of error detection during skilled performance

**DOI:** 10.1101/484584

**Authors:** Gabriela Bury, Marta García Huesca, Joydeep Bhattacharya, María Herrojo Ruiz

**Author notes:** **Corresponding author:** María Herrojo Ruiz,. Address: Department of Psychology, Goldsmiths, University of London. Lewisham Way, New Cross, London SE14 6NW (UK).

## Abstract

Behavioral adaptations during performance rely on predicting and evaluating the consequences of our actions through action monitoring. Previous studies revealed that proprioceptive and exteroceptive signals contribute to error-monitoring processes, which are implemented in the posterior medial frontal cortex. Interestingly, errors also trigger changes in autonomic nervous system activity such as pupil dilation or heartbeat deceleration. Yet, the contribution of implicit interoceptive signals of bodily states to error-monitoring during ongoing performance has been overlooked.

This study investigated whether cardiovascular interoceptive signals influence the neural correlates of error processing during performance, with an emphasis on the early stages of error processing. We recorded musicians’ electroencephalography and electrocardiogram signals during the performance of highly-trained music pieces. Previous event-related potential (ERP) studies revealed that pitch errors during skilled musical performance are preceded by an error detection signal, the pre-error-negativity (preERN), and followed by a later error positivity (PE). In this study, by combining ERP, source localization and multivariate pattern classification analysis, we found that the error-minus-correct ERP waveform had an enhanced amplitude within 40-100 ms following errors in the systolic period of the cardiac cycle. This component could be decoded from single-trials, was dissociated from the preERN and PE, and stemmed from the inferior parietal cortex, which is a region implicated in cardiac autonomic regulation. In addition, the phase of the cardiac cycle influenced behavioral alterations resulting from errors, with a smaller post-error slowing and less perturbed velocity in keystrokes following pitch errors in the systole relative to the diastole phase of the cardiac cycle. Lastly, changes in the heart rate anticipated the upcoming occurrence of errors. This study provides the first evidence of preconscious visceral information modulating neural and behavioral responses related to early error monitoring during skilled performance.

## Introduction

Detection and evaluation of errors is central to the acquisition of complex motor skills, such as those needed in dance or music performance (Wolpert and Kawato, 1998; Palmer, 2006). Over the last decades, error-monitoring processes have received broad attention in the neuroscientific community (Gehring et al, 1993; Falkenstein et al, 2000; Ullsperger et al., 2014). Electroencephalography (EEG) has been the most widely used method to investigate the neural correlates of error-monitoring processes, as its high-temporal resolution in the order of milliseconds makes it particularly suitable for tracking the fast brain dynamics associated with error processing (Delorme et al., 2002). Changes in neural activity around error commission have been traditionally assessed by averaging the EEG signals time-locked to errors, thus rendering the classical event-related potential (ERP) responses, the early (80-100 ms after error commission) error-related negativity (ERN, Gehring et al, 1993), and a later error positivity (PE, Falkenstein et al, 2000).

The ERN is generated in the posterior medial frontal cortex (pMFC), anterior cingulate cortex (ACC), and pre/suplementary motor area (pre-/SMA) (Kiehl et al., 2000; Miltner et al. 2003; Bonini et al., 2014). This ERP component is considered to reflect a mismatch between the ongoing expectation of action outcomes and the actual outcomes (Hyman et al., 2017; Fu et al., 2019), a process which has been shown to involve the pMFC and ACC (Rushworth and Behrens, 2008; Fu et al., 2019). Following extensive training, for instance during skilled musical performance, early error detection in the pMFC/ACC precedes error commission by 50-100ms (pre-error-negativity, preERN; Maidhof et al., 2009, Ruiz et al., 2009; Strübing et al., 2012).

Oscillatory analysis of the preERN suggests that the theta (4-7 Hz) and beta (13-30 Hz) oscillations in the pMFC primarily contribute to the ERP modulation, and connectivity analyses have shown an increase in beta band coupling between the pMFC and the lateral prefrontal cortex (lPFC) in anticipation of an error (Ruiz et al., 2011). Further, in that study, this enhanced connectivity was associated with larger corrective behavioral adjustments.

Whereas early error detection occurs in the pMFC, later error evaluation – linked to the PE component – also extends to subcortical regions. The PE consists of an earlier frontocentral and a later centroparietal deflection, which peak at 200 and 300 ms following errors, respectively. The later deflection, in particular, reflects an accumulation of evidence or decision confidence that an error has occurred (Orr and Carrasco, 2010; Boldt and Yeung, 2015). Interestingly, the PE is generated in the rostral ACC and the insular cortex (Herrmann et al., 2004; Dhar et al., 2011). Thus, the neural signatures of error detection and evaluation rely on an extended cortico-subcortical network beyond the pMFC (Ullsperger et al., 2014).

The anterior insular cortex is a key region for interoception, defined as the sense of the internal physiological state of the body (Craig, 2009; Barrett and Simmons, 2015). Accordingly, the involvement of this brain region during error processing aligns well with the reported changes in the ongoing autonomic nervous system, such as heartbeat deceleration and enhanced pupil dilation, following errors (Hajcak et al., 2003; Wessel et al., 2011, Bastin et al., 2017). Further, the ventral part of the ACC is a key recipient of visceral information from the internal organs (Stevens et al., 2011; Critchley and Harrison, 2013). Overall, these findings suggest that error commission may be associated with interoceptive processing, yet we do not fully understand how the interoceptive signals of bodily states contribute to error-monitoring during ongoing performance. This is surprising given the relevance of interoceptive information for the modulation of perceptual, affective and cognitive processes (Critchey and Harrison, 2013; Park et al., 2014). Recent work has further shown that changes in interoceptive states (heart rate deceleration) following incorrect responses might provide internal feedback about performance accuracy (Lukowska et al., 2018). Also following errors, the skin-conductance response – a measure of the autonomic nervous system – correlates with the PE amplitude and the degree of post-error slowing (Hacjak et al., 2003). What remains unclear is whether interoceptive cues influence error-monitoring processes during earlier stages. New theoretical accounts suggest that interoceptive processing is not limited to monitoring the physiological state of the body but is also crucially involved in generating future predictions (expectations) about bodily states (Seth, 2013; Barrett and Simons, 2015). Accordingly, early interoceptive cues could shape ongoing error-monitoring to influence the predicted future bodily states associated with error commission.

The present study tested the general hypothesis that early detection of errors during skilled motor performance is influenced by implicit cardiovascular interoceptive information. Skilled performance of motor tasks, such as music performance, sports, or dance demands extensive training and perfect tuning of the action-monitoring system to the extent that potential errors, which might otherwise interact with the goals, must be anticipated (predicted; Bernstein, 1967; Wolpert et al., 1995). A highly-trained music performance presents thus a suitable paradigm for the study of the brain’s predictive mechanisms of error detection. We predicted that cardiovascular interoceptive information could shape error processing during skilled music performance, given the recent evidence that interoceptive processing can modulate error responses (Bastin et al., 2018; Lukowska et al., 2018), and the studies revealing that expert musicians (singers) have higher interoceptive accuracy likely due to the intensive training (Schirmer-Mokwa et al., 2015),

Here, we recorded EEG and electrocardiogram (ECG) of skilled pianists during performance of highly-trained musical excerpts. There are two dominant approaches to studying the relationship between neural and cardiovascular responses during cognition. First, analysis of the heartbeat-evoked potential (HEP) – neural responses locked to heartbeats – has been used to demonstrate a role of visceral information in bodily consciousness, conscious perception and emotional processing (Pollatos and Schandry 2004; Luft and Bhattacharya, 2015; Babo-Rebelo et al., 2016a and 2016b; Park et al., 2016). This approach is typically followed in studies that do not measure concurrent motor responses, which could overlap with the HEP modulation. A second approach consists of measuring ongoing fluctuations of the cardiac cycle; these fluctuations have been shown to influence perceptual and sensory processing (Garfinkel et al., 2014; Azevedo et al., 2017; Al et al., 2018). The cardiac cycle extends from the contraction of the atria to the ventricular relaxation (Lacey and Lacey, 1978: Gray et al., 2009). It can be characterized by two phases: the systole, when the ventricles contract and blood is pumped into the arteries, and the diastole, when the ventricles relax and are filled with blood. Numerous studies have explored the effect of stimuli presented during the different phases of the cardiac cycle on neurophysiological and psychological processes, demonstrating that processing of fearful or painful stimuli, i.e., stimuli associated with enhanced emotional arousal, is more efficient during cardiac systole (Edwards et al., 2001; Garfinkel et al., 2014; Azevedo et al., 2017). Notably, this second approach has been preferred in studies measuring the effect of cardiac activity on motor behavior. These studies have demonstrated that the cardiac phase can have an effect on motor task performance, either through a modulation of motor preparation or sensory processing (in stimulus-response paradigms; Weisz and Adam, 1996; Edwards et al., 2007). For example, response to a stimulus presented during the cardiac systole can be slower, possibly due to an increase in blood pressure when the arterial baroreceptors are naturally stimulated by the arrival of the pulse pressure wave during systole (Edwards et al., 2007).

In this study, we adopted this second approach as it is well-suited to capturing cardiovascular effects on motor behavior, such as during music performance. Accordingly, we undertook a cardiac-cycle fluctuation-based analysis to assess whether the alignment of performance errors to different phases of the cardiac cycle biases neural and behavioral responses during error processing. Rhythmic feedback of cardiac and baroreceptor activity constitutes one important channel of brain-body communication (Gray et al., 2009), with baroreceptors showing maximal phasic activity during the systolic period, thereby enabling the representation of cardiovascular arousal within viscerosensory brain regions (Azevedo et al., 2017). Accordingly, our specific hypotheses were as follows: (I) ERP responses to errors were expected to be modulated in amplitude depending on the latency of the error key presses within the cardiac cycle. (II) Behavioral alterations to errors were predicted to be enhanced during the cardiac systole, corresponding with the more salient processing of negative arousal (painful, fearful) stimuli during systole (Edwards et al., 2001; Garfinkel et al., 2014; Azevedo et al., 2017). (III) The ERP modulation by the cardiac cycle should stem from brain regions previously associated with processing of cardiovascular afferent signals, such as the ACC, somatosensory cortex or the inferior parietal cortex (Lutz et al., 2009; Park et al., 2014; Barrett and Simons, 2015). Finally, (IV) ongoing changes in the autonomic nervous system, such as heart-rate acceleration or deceleration, were expected to be precursors of the early error components. To test these hypotheses and dissociate between the neural responses to pitch errors during the different phases (systole, diastole) of the cardiac cycle, we used different methods, including ERP, source localization and multivariate pattern classification techniques.

In addition to the cardiac-cycle fluctuation-based analysis, we further aimed to replicate the previous results of the preERN and PE preceding and following pitch errors, as well as the behavioral effects surrounding errors (pre- and post-error slowing and reduced keystroke velocity in pitch errors, Ruiz et al., 2009; Maidhof et al., 2009; Palmer et al., 2012). This additional goal was motivated by the recent call for replication of small scale neuroimaging studies (Button et al., 2013; Open Science Collaboration, 2015).

## Materials and Methods

### Participants

Sample size considerations were based on prior data from our study using this paradigm (Ruiz et al., 2009), which we used to estimate the minimum sample size for a statistical power of 0.95, with an alpha of 0.05, using the MATLAB function sampsizepwr (https://de.mathworks.com/help/stats/sampsizepwr.html). Using the preERN and PE data from the 19 pianists that participated in that study, we obtained a minimum sample size of 17. In addition, the non-parametric effect sizes of the previously reported preERN and PE were 0.875 and 0.812, respectively (non-parametric effect size estimator, Grissom and Kim; 2012).

Eighteen right-handed healthy pianists (10 females; mean age = 26.6, s.d. = 2.7 years; Handedness assessed with the Edinburgh inventory, Oldfield, 1971) participated in the present study. They were professional pianists or current students at a London music conservatoire, and had self-reported normal hearing. They were all practicing musicians and had accumulated more than 10,000 hours of practice. All participants gave written informed consent, and the study protocol was approved by the local ethics committee at the Department of Psychology, Goldsmiths, University of London. Participants received a monetary remuneration for participating in the study and memorising the musical material prior to their participation. One participant was excluded from the analysis due to poor quality of the raw EEG signal, leaving 17 participants for the analysis.

### Stimulus Materials

Stimuli were four musical sequences selected from the material used in Ruiz et al, (2009). In brief, the sequences were right-hand excerpts of Preludes V and VI of the Well-Tempered Clavier by J. S. Bach. These pieces were chosen because of their homogeneity in terms of the duration of notes (sixteenth notes) and regular (isochronous) time interval between consecutive notes. The length of these excerpts was 200, 201, 202 and 185 notes, respectively. As in Ruiz et al. (2009), participants were instructed to perform at a fast tempo (metronome 120 bpm for quarter note in excerpts from prelude V and 160 bpm for triplet of sixteenth notes in excerpts from prelude VI) to induce errors. The tempo indications thus required a target inter-onset-interval, IOI, of 125ms, between sixteenth notes. The duration of each piece was around 25 s. Participants were instructed to rehearse and memorize the pieces at the correct tempo while not looking at their hand and finger movements before coming to the experimental session. Prior to EEG and performance recording, we verified that pianists managed to play all pieces from memory without looking at their hands.

### Experimental Design

Participants were seated comfortably on an adjustable piano stool in front of a digital piano (Yamaha Digital Piano P-255, London, United Kingdom) in a dimly lit room. They were instructed, via a PC monitor, to perform each piece from beginning to end at the instructed tempo and without stopping to correct errors. They were also instructed to avoid visually tracking their finger movements by fixating on a central cross displayed on the PC monitor. The emphasis was placed on maintaining accurate timing (see below) and playing the correct notes. All participants were naïve to the purpose of the study.

The experiment consisted of 60 trials, separated into two 30-trial blocks of approximately 20 min each. There were two practice trials at the beginning of each block for familiarization. We used a synchronization-continuation paradigm with the following timeline for each trial. First, participants pressed a designated key with their left-hand index finger when ready to begin a trial, which was followed by a silent time-interval between 0-1000ms (jitter). Next, an image with the first two bars of the music score was displayed for 4000 ms, indicating which of the four excerpts, randomly selected, had to be played. While the visual cue (score) was shown, a metronome started pacing the tempo corresponding to the piece: four metronome beats for scores at 120 bpm (at regular intervals of 500 ms), five metronome beats for ones at 160 bpm (at 375ms-intervals). Participants had to internally entrain their timing to the metronome clicks and get ready to start playing after the last metronome beat. In every trial, the last metronome beat fell at 4050 ms after the initial presentation of the music score, and 50 ms after a green ellipse was shown on the monitor to indicate that performance was being recorded. Participants were instructed to play while the green ellipse was on display and to stop playing when the ellipse turned red (approximately after 27 s).

### EEG, ECG and MIDI recording

EEG and ECG signals were recorded using a 64-channel (extended international 10–20 system) EEG system (ActiveTwo, BioSemi Inc.) placed in an electromagnetically shielded room. During the recording, the data was high-pass filtered at 0.16 Hz. The vertical and horizontal eye-movements (EOG) were monitored by electrodes placed above and below the right eye and from the outer canthi of both eyes, respectively. Additional external electrodes were placed on both left and right mastoids to use as initial references upon importing the data in the analysis software (data were subsequently re-referenced to common average reference, see below). The ECG was recorded using two external channels with a bipolar ECG lead II configuration (the negative electrode was placed on the chest bellow the right collar bone, and the positive electrode was placed on the left leg above the hip bone). The sampling frequency was 512 Hz. Performance was additionally recorded as MIDI (Musical Instrument Digital Interface) files using the software Visual Basic and a standard MIDI sequencer program on a PC with Windows XP software (compatible with Visual Basic and the MIDI sequencer libraries we used). The MIDI controller was set to the default linear velocity mapping (range of keystroke velocity 0-127). To run the behavioral paradigm and record the MIDI data, we used a modified version of the custom-written code for Visual Basic that was used in similar paradigms in our previous studies (e.g. Herrojo Ruiz et al., 2009). This program was also used to send synchronization signals in the form of transistor–transistor logic (TTL) pulses – corresponding with onsets of visual stimuli, key presses and metronome beats – to the EEG/ECG acquisition PC.

### EEG Data Preprocessing

We used MATLAB (The MathWorks, Inc., MA, USA) and the FieldTrip toolbox (Oostenveld et al. 2011) for visualization, filtering and independent component analysis (ICA; runica). The EEG data were highpass-filtered at 0.5 Hz (Hamming windowed sinc FIR filter, 3380 points) and notch-filtered at 50 Hz (847 point). ICA was used to identify artifact componets related to eye blinks, eye movements and, crucially, the cardiac-field artifact (CFA). Inspection of the IC components to find the CFA was performed using the procedure suggested in the Fieldtrip toolbox (http://www.fieldtriptoolbox.org/example/use_independent_component_analysis_ica_to_remove_ecg_artifacts).

Following this approach, the time and the frequency data of the ECG signal were used to first detect the QRS-complex, which is the main spike seen on the ECG signal corresponding with the depolarization of the heart ventricles, corresponding with the cardiac systole. The largest peak in the complex is the R wave peak. Fieldtrip uses an efficient peak detection algorithm based on thresholding the z-transformed values of the preprocessed raw data (ECG signals for detection of the R-wave peak). Next, the time course of the QRS-complex was correlated with the IC time series. We typically found one or two ICs related to the CFA, which were subsequently removed. After IC inspection, we used the EEGLAB toolbox (Delorme and Makeig, 2004) to interpolate missing or noisy channels using spherical interpolation. Next, the data were re-referenced to a common average reference.

In the analysis of epoched data, we performed a final inspection of the epoched data with the EEGLAB toolbox and applied an automated rejection of noisy epochs exceeding a voltage threshold of ±100 μV.

### ECG data Preprocessing

The analysis of the ECG signal with Fieldtrip rendered the time course of the heartbeat and associated QRS complex. We then dissociated between different types of R-peaks, depending on the temporal difference between the R-peak and the subsequent keystroke (pitch error or correct note). Specifically, we selected R-peaks which were followed by a keystroke either (i) within 50-350 ms, corresponding to the systolic period of the cardiac cycle and the T waveform (peaking at 233 [standard error of the mean or s.e.m. 8] ms after the R-peak on average); or (ii) between 400 ms and the beginning of the next R-peak (693 [15] ms on average), corresponding to the diastole phase of the cardiac cycle (Figure 1). The systolic period, in which the blood is pumped out of the heart, is the window of maximal representation of baroreceptor afferent activity (Fadel et al., 2003) and when cardiac cycles effects on perception and cognition are typically observed (Motyka et al., 2007; Gray et al., 2012; Garfinkel et al., 2014). The diastole is the remaining period of the cardiac cycle during which afferent activity is quiet. From hereafter, events (e.g. keystrokes) in each of these periods of the cardiac cycle will be labelled as pertaining to bin1 (systole) or bin2 (diastole), respectively.

**Figure 1.**
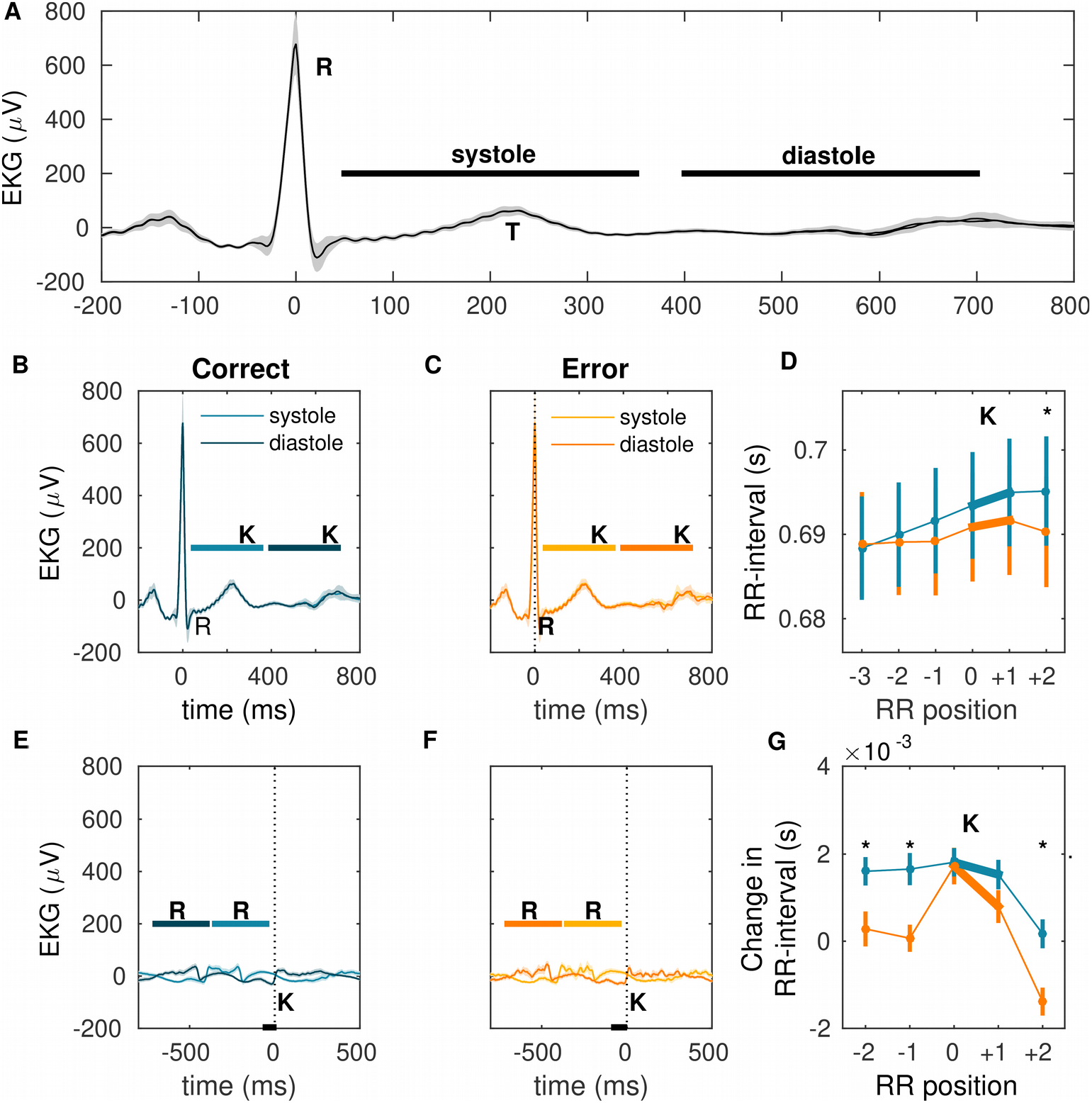
Electrocardiogram (ECG). Grand-average across subjects. Shaded areas show ± 1 s.e.m. around the mean. **(A)** Scheme representing the systolic and diastolic phases of the cardiac cycle in the ECG used for the analyses in our experiment. The R-wave peak, as well as the T-wave peak are also highlighted by letters R and T, respectively. **(B)** Analysis of ECG locked to the R-peak for correct notes falling within the systolic (light blue) or diastolic (dark blue) phase of the cardiac cycle revealed no significant differences (p > 0.05). The blue horizontal likes denoted by “K” highlight the temporal interval in which the keystroke fell for each phase of the cardiac cycle. The interval was 50-350 ms for the systolic and from 400 ms until the next R-peak (693 ms on average, standard error of the mean or s.e.m. 15 ms) for the diastolic phase. The vertical line denoted by ‘R’ indicates the onset of the R peak. **(C)** Same as (A) but in error trials. No significant differences between the R-peak-locked ECG in bin1 (systole, light orange) and bin2 (diastole, dark orange) trials was found (p > 0.05). **(D)** Heart-rate changes around pitch errors (in orange) compared to correct key presses (in blue) measured as RR-interval (ms) and shown as mean (in seconds) and s.e.m. bars. The thicker line segments from RR-interval position 0 to 1 highlight the time interval in which the error or correct keys were pressed (also denoted by “K”). The RR-interval differed significantly between error and correct key presses exclusively at position +2K following keystrokes, due to a relative faster heart rate following errors (p < 0.05, FDR-corrected, PS_dep_ = 0.94). **(E)** ECG locked to the keystroke following R-peaks and separately for bins 1 and 2 in correct trials. Time 0 ms marks the correct note onset, denoted by the vertical line (labeled “K”). The shifted ECG waveforms in bin1 and bin2 were significantly different within [−53, −8] ms (black bottom line; p < 0.05, FDR-corrected; PS_dep_ = 0.875). The horizontal lines in light and dark blue denoted by ‘R’ indicate the temporal intervals of the R-peak preceding the keystrokes in each bin. **(F)** Same as (E) but for error trials. The shifted ECG waveforms locked to pitch errors differed significantly between bin1 and bin2 trials also in the pre-keystroke interval, similarly to the results in correct trials, within [−74, −18] ms (p < 0.05, FDR-corrected; PS_dep_ = 0.818). **(G)** Same as (D) but for differences between consecutive RR-intervals (e.g. RR-interval at position 0, before the keystroke, relative to the previous RR-interval). The RR-interval change prior to error keystrokes was smaller than in error trials (−2, −1, positions) as well as following the keystrokes at the +2 RR-interval position (p < 0.05, FDR-corrected; PS_dep_ in range 0.76 – 0.85). The pronounced drop in RR-interval change following errors relative to correct notes indicated that pianists’ heart rate speeded up after a pitch error, yet slowed down prior to error commision (positive change at position 0 relative to the previous one −1 in the RR-interval; p < 0.05, FDR-corrected; PS_dep_ = 0.84).

### Performance analysis and performance-related modulation of the heartbeat

Consistent with Ruiz et al., (2009), we analyzed the MIDI files to detect pitch errors during piano performance. In addition, we extracted information corresponding to the timing of note onsets, the inter-onset-interval (IOI, ms), i.e. the time between consecutive keystrokes, and the keystroke velocity, related to the loudness. As in our previous studies (Ruiz et al. 2009, 2011), we exclusively selected “isolated” pitch errors and correct notes as events for the EEG analysis. Isolated pitch errors and correct notes refer to events surrounded by correct pitch notes for at least two preceding and following keystrokes. We also constrained the minimal and maximal IOI prior to and post-keystroke to 100 and 300 ms, respectively. General performance was assessed in terms of average timing (mean IOI, mIOI, in ms), temporal variability (coefficient of variation of IOI, cvIOI), average MIDI keystroke velocity (mKvel, related to loudness), and error rates.

To assess changes in the heart rate (HR) preceding and following error commission (e.g. Bastin et al., 2017), relative to performance of correct note onsets, we used the time series of R-peaks and inter-beat intervals (termed RR-interval hereafter, ms).

### Event-related potentials: Keystroke-locked preERN and PE analysis

Before analysing the influence of cardiac interoceptive signals on error-processing, we first tried to replicate previous results reporting the emergence of a preERN ([−80, −30] ms) and PE ([150, 250] ms) ERP components triggered by pitch errors during piano performance (Ruiz et al., 2009; Maidhof et al., 2009). To that aim, analysis of keystroke-locked data focused on isolated pitch error and correct note events (see previous section). Epochs were extracted within [−1000, 1000] ms around the keystroke. After EEG preprocessing and artifact removal, the average number of trials available for this analysis was 170 (s.e.m.15) errors and 1600 (200) correct notes. Because pitch errors are typically slower than correct key presses, and the difference in latencies could lead to a modulation of the ERP waveforms (see Maidhof et al. 2013), we performed a control analysis by selecting a subset of the pitch error events with comparable timing to that of isolated correct events. Specifically, for each subject, we selected pitch error events with an IOI falling within the subject-specific range of IOIs for isolated correct events. This resulted in a group-average of 73 (9) pitch error trials available (no difference in IOI between this subset of error trials and correct events, 126 (1) ms and 125 (0.1) ms, respectively, p = 0.3250). After we controlled for temporal differences between types of events, this analysis replicated the previous ERP findings (see below). Accordingly, the main analyses throughout the manuscript focused on the total set of pitch error trials as a larger number of trials was thought to improve the signal-to-noise ratio.

The analysis (see *Statistical Analysis*) was carried out within [−200, 300] ms as this interval contains the previously reported preERN ([−80, −30] ms) and PE ([150, 250] ms) as well as a potential effect preceding the pre-error keystroke (Maidhof et al., 2013). The ERP data were corrected with a baseline reference level from −300 to −150 ms prior to the keystroke (Ruiz et al., 2009).

The ERP waveforms were additionally assessed by comparing trials in which the keystroke coincided with the systolic (bin1) or diastolic (bin2) phase of the cardiac cycle. The number of available epochs of each type for this analysis were: 74 (6) errors in bin1 and 69 (7) in bin2; 629 (70) correct events in bin1 and 580 (70) in bin2. There were no significant differences between bins in the number of available epochs (p > 0.05 for error and correct events).

### Source Reconstruction

Between-conditions differences at the source level were assessed using L2-norm minimum-norm estimates (MNEs, Hämäläinen and Ilmoniemi, 1994; Dale et al., 2000) and a standard brain T1-weighted MRI template (Colin27 brain in MNI152 space). MRI segmentation was performed with a FieldTrip pipeline (http://www.fieldtriptoolbox.org/workshop/ohbm2018) to generate a volumetric mesh as a head model with five compartments: scalp, skull, cerebrospinal fluid (csf), gray and white matter. Co-registration of the MRI and EEG coordinate system was then carried out using the anterior and posterior commissure as the anatomical landmarks. Next, the FieldTrip-SimBio pipeline was implemented to calculate EEG forward solutions using the finite element method (FEM; Vorwerk et al., 2018). Next, inverse solutions were computed using the L2-norm MNE method, as implemented in the “FieldTrip” software (minimum-norm estimate, based on Dale et al., 2000; Lin et al., 2004). MNE sources were estimated for each grid point in the window of the significant ERP effect, the individual source solutions were interpolated to a template MNI mesh. The noise-covariance matrix was estimated for each participant using a time window preceding the error and correct events, within [−0.7, −0.3] s. Scaling of the noise-covariance matrix was performed with regularization parameter λ = 0.1.

### Multivariate Pattern Classification Analysis (MVPA)

We first used MVPA (Duda et al., 2000; Haxby et al., 2001) to assess whether error and correct trials could be classified using the heart rate (HR) information from the three preceding interbeat intervals (three features).

In addition, for the EEG analysis, MVPA was used to determine whether the phase of the cardiac cycle (systole vs diastole) could be decoded from the neural responses to errors on a trial-by-trial basis (Duda et al., 2000; Haxby et al., 2001). In this analysis we used a subset of the data from each participant, to match the number of error trials from systolic and diastolic phases. The features selected for MVPA were the ERP amplitude values in the set of 64 electrodes. MVPA was performed in the range [−150, 300] ms – similar to the interval used for the standard ERP analysis. Each data epoch was normalized using this interval (full-epoch-length normalization: The mean from the full epoch is subtracted, instead of removing the pre-event baseline, a method widely used when normalizing single trials; see Grandchamp and Delorme, 2011). The range −150 to 300ms was divided into 50 adjacent windows of 10-ms width.

For the HR-MVPA, a support vector machine (SVM, a MATLAB library by Chang and Lin, 2011) with a radial basis kernel function was trained to distinguish between error and correct trials. For the EEG-MVPA, however, a SVM with a radial basis kernel function was trained separately in each time window to distinguish between trial classes (keystroke in the systolic or diastolic period).

In all MVPA cases, we used 10-fold leave-one-out cross-validation to estimate the validity of the SVM model. A participant-based classification was performed first, and subsequently, we estimated the population decoding accuracy by averaging across participants.

### Statistical Analysis

Statistical analysis of group-level behavioral data was performed using pair-wise permutation tests (Good, 2005), using the difference between condition means as the test statistic. Whenever multiple comparisons were performed, we applied an adaptive two-stage linear step-up procedure to control the false discovery rate (FDR) at level *q* = 0.05 (Benjamini et al., 2006; termed *p* < 0.05, FDR-corrected hereafter).

The group-level statistical inference on the difference between error and correct trials in keystroke-locked ERPs was conducted using permutation tests with a cluster-based threshold correction to control the family-wise error (FWE) at level 0.05 (dependent samples t-test, 1000 iterations; Maris and Oostenveld, 2007). Experimental cluster-based test statistics being in the 2.5^th^ and 97.5^th^ percentiles of the permutation distribution were considered significant (two-tailed test, *p* < 0.025). Using this procedure, the assessment of statistical differences in the keystroke-locked ERP was performed within [−100 ms, 300 ms]. To assess the influence of cardiac information on neural responses we compared the keystroke-locked error-minus-correct ERPs between bin1 (systole) and bin2 (diastole) epochs. Statistical analysis for this comparison was also performed within [−100, 300] ms.

The statistical analysis was complemented with a nonparametric effect size estimator, the probability of superiority for dependent samples or PS_dep_ (Grissom and Kim; 2012). PS_dep_ is an estimation of the probability (maximum 1) that in a randomly sampled pair of matched values *i* (from same individual), the value from Condition B will be greater than the value from Condition A: PS_dep_ = Pr (XB_i_ > XA_i_). Throughout the paper, this index will be provided in association with each pairwise permutation test (excluding the cluster-permutation test on spatiotemporal features, for which PS_dep_ is not defined).

Differences between conditions in source reconstruction were also assessed at the group level using the cluster-based permutation test (Maris and Oostenveld, 2007) As in the case of statistics at the channel level, correction for multiple comparisons at the source level was performed by controlling the FWE at level 0.05 (and alpha level 0.025, two-tailed test).

Finally, for the MVPA, we first tested whether the classification accuracy in each participant was significant at the participant level. To that aim, the null distribution of accuracy in each participant was estimated by performing the MVPA 500 times after randomly shuffling the class labels in the data. At each time window (EEG-MVPA) or at once (HR-MVPA) we calculated the p-values as the frequencies of permutation accuracies that are greater than or equal to the experimental decoding accuracy. Note that this approach to assessing statistical significance on the subject level (see also Ruiz et al., 2014) is preferred than comparing empirical classification accuracies against a theoretical chance level of 50% for two-class classification. The reason is that the empirical chance level in two-class classification problems prominently deviates from 50% for small sample sizes (small number of trials; Combrisson and Jerbi, 2015).

For the EEG-MVPA, the single-subject MVPA thus aimed to complement the cardiac cycle-based ERP group results by revealing in individual participants the effect of the phase of the cardiac cycle on the modulation of neural responses to errors. Next, statistical assessment with a permutation test was performed on the group level to localize the time windows showing an effect of above-chance decoding accuracy across the pianist population. The population chance level used in this analysis was the mean across individual permutation-based chance levels, extracted as the mean of the null distribution of decoding accuracies in each participant. Of note, however, for MVPA single-subject statistical assessment might be preferred over group-level statistical analysis when using the mean accuracy as test statistic, as statistical inference on the group level using this test statistic is not optimal to describe whether effects related to information content are how “typical” in the population (that is, whether they are present in the majority of the population; Allefeld et al., 2016).

## Results

### Behavioral Results

The details of the behavioral data are shown in Table 1. All data are provided as mean and s.e.m. Our participants played the musical excerpts at the instructed tempo (average time between consecutive notes, or IOI, was 125 [s.e.m. 0.1] ms), when excluding pitch errors in the analysis. The percentage of isolated pitch errors was 1.4% (0.2%), and the percentage of total pitch errors was 3.7% (1%). Pitch errors were distributed across all positions of each excerpt, indicating that they were not linked to an isolated part of the sequence. Isolated pitch errors fell into two categories, with pitch replacement being the most frequent type of error (rate of 0.68 [0.05]), followed by pitch omission (one tone was skipped, 0.32 [0.05]). This distribution of larger rate of isolated pitch errors of the pitch replacement type was consistent across 11 participants, whereas in only two participants there were more omissions than pitch replacements. A similar rate of both types of isolated pitch errors was found in four participants.

**Table 1.** Performance data expressed as mean (s.e.m.). IOI (ms) across different keystroke positions relative to the target keystroke is denoted by “K” (positions +1, +2, +3, −1, −2, −3).

|  |  |
| --- | --- |
| Percentage of total pitch errors | 3.7% (1%) |
| Percentage of isolated pitch errors | 1.4% (0.2%) |
| Percentage of isolated pitch errors: pitch replacement | 0.95% (0.09%) |
| Percentage of isolated pitch errors: omission | 0.45% (0.09%) |
| Number of total pitch errors | 400 (90) |
| Number of isolated errors | 153 (16) |
| Number of isolated pitch errors: pitch replacement | 104 (11) |
| Number of isolated pitch errors: omission | 49 (11) |
| Number of total notes played | 10600 (380) |
| Mean IOI of correct pitch keystrokes (ms) | 125 (0.1) |
| Mean IOI of isolated pitch errors (ms) | 169 (8) |
| Overall loudness: Correct pitch | 66 (1) |
| Overall loudness: Pitch errors | 64 (1) |
| Mean IOI at +K following isolated pitch errors (ms) | 149 (8) |
| Mean IOI at +2K following isolated pitch errors (ms) | 147 (7) |
| Mean IOI at +3K following isolated pitch errors (ms) | 143 (6) |
| Mean IOI at -K preceding isolated pitch errors (ms) | 134 (6) |
| Mean IOI at -2K preceding isolated pitch errors (ms) | 131 (6) |

In terms of timing, isolated pitch errors had a slower tempo (larger IOI relative to previous keystroke) than correct key presses (p = 0.002, p < 0.05, FDR-corrected, PS_dep_ = 1; Table 1). With regard to MIDI velocity, pitch errors had a significantly smaller value than correct key presses (64 [1] and 66 [1], respectively, p = 0.006, p < 0.05, FDR-corrected, PS_dep_ = 0.76). Timing and keystroke velocity changes at the error were consistent across the two types of isolated pitch errors (pitch replacement: IOI = 185 [30] ms, Kvel = 64 [1]; omissions: IOI = 174 [20] ms, Kvel = 63 [3]; no significant differences between types, p > 0.05).

Timing and keystroke velocity data at keystrokes surrounding pitch errors were consistent with post-error adaptation effects but also with pre-error alterations in performance. Specifically, pitch errors were followed by significantly slower (delayed) keystrokes in the subsequent +K, +2K, +3K, +4K, +5K and +6K events and preceded by a slower keystroke at −K relative to correct events at those same positions (p < 0.05, FDR-corrected in all cases; PS_dep_ in the range 0.73-0.94; nonsignificant difference at −2K and +7K, p > 0.05; Figure 2). The slowing of IOI following pitch errors relative to correct key presses was, however, less pronounced than the slowing of IOI at the errors (p < 0.05, FDR-corrected in all cases +K to +6K; PS_dep_ in the range 0.71-0.87). A significantly reduced MIDI velocity was also observed at the previous and subsequent keystrokes relative to the correct notes in those positions (65 [1] at −K and +K relative to errors; 66 [1] at events surrounding correct note onsets, p < 0.05, FDR-corrected, PS_dep_ 0.94 and 0.59, respectively). Consequently, behavioral alterations in performance were most pronounced at pitch errors, yet extended to the keystroke preceding as well as several keystrokes following pitch errors.

**Figure 2.**
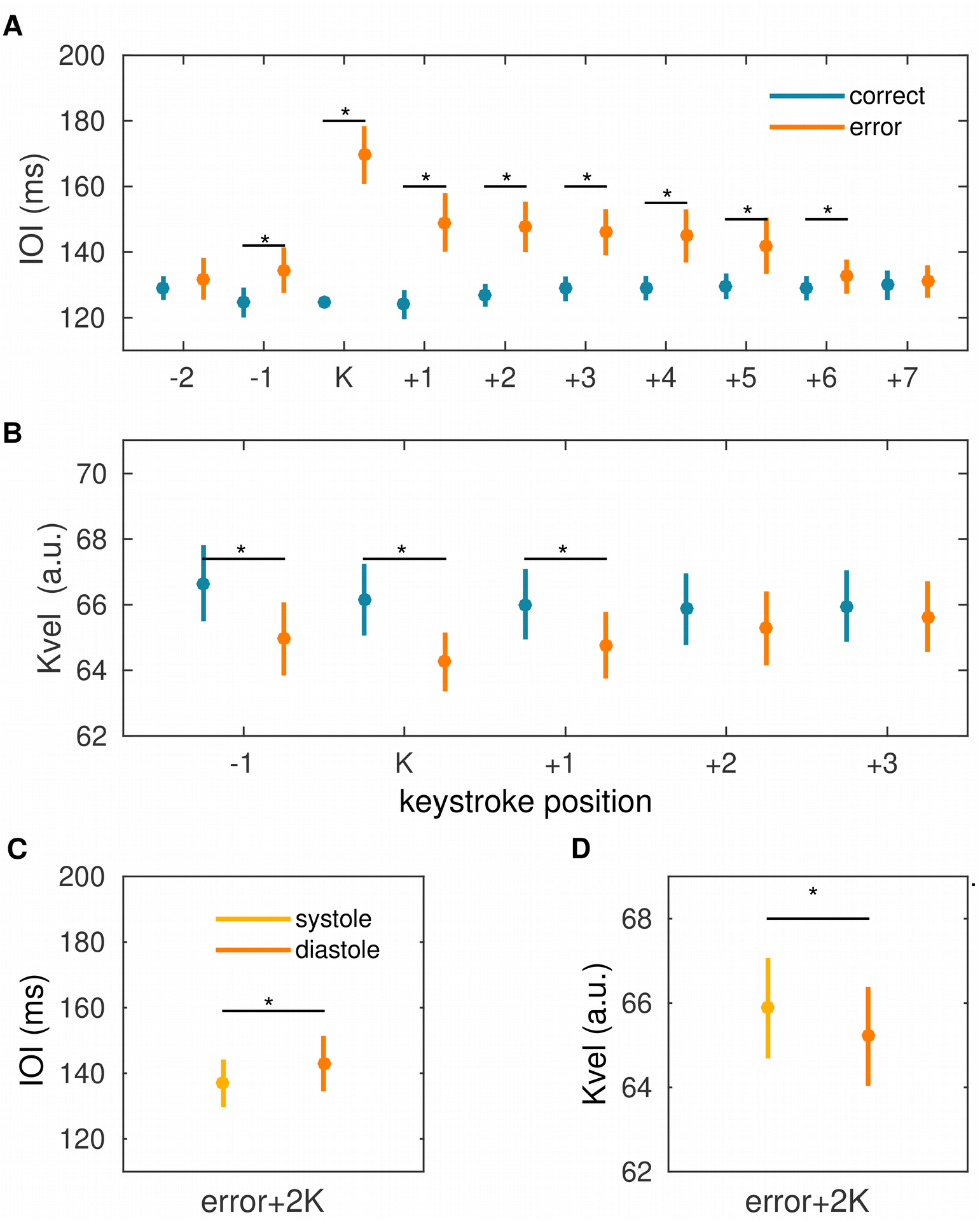
Behavioral data. Data are presented as mean and s.e.m. bars. **(A)** Mean inter-onset-interval, IOI (ms), across different keystroke positions relative to the target keystroke (pitch errors, in orange; correct note, in blue), denoted by “K”. The mean timing of keystrokes before and after pitch errors was significantly slower than the average tempo of keystrokes surrounding correct key presses at the same position (positions −1, +1, +2, +3, +4, +5, +6; p < 0.05, FDR-corrected; PS_dep_ in range 0.73-0.94; significant differences denoted by the horizontal black lines with an asterisk). Pitch errors were also slower than correct keystrokes (p < 0.05, FDR-corrected; PS_dep_ = 1). **(B)** The mean keystroke velocity, associated with the loudness, at pitch errors as well as preceding and following errors was significantly reduced compared to the keystroke velocity of correct notes at the same position (positions −1, target keystroke, +1; p < 0.05, FDR-corrected; PS_dep_ was 0.94, 0.76, 0.59, respectively). **(C)** The slowing of performance timing at pitch errors and at +2K following errors was significantly smaller when the pitch errors coincided with the systolic (light orange), relative to the diastole (dark orange), period of the cardiac cycle (p < 0.05, FDR-corrected; PS_dep_ = 0.77 and 0.69, respectively).

The subset of pitch errors selected to match correct events in timing (IOI, ms; see Methods), which were subsequently used in a control ERP analysis, exhibited significant post-error slowing effects. Specifically, the mean IOI of the keystrokes following the pitch error at +K, +2K and +3K positions was 136 (4) ms, 137 (5) ms, and 137 (4) ms, respectively, thus being larger than the average IOI, 126 (1) ms, of the subset of pitch errors (p < 0.05, FDR-corrected). The keystroke velocity was also reduced at position +K following the pitch error (65 [1], which was smaller than in correct events: p < 0.05, FDR-corrected). Therefore, the general effects of post-error slowing in tempo and reduced loudness were maintained after we controlled for the timing of the pitch errors to match the values of the correct events.

### Effects of behavioral performance on the heart rate

On average, the cardiac interbeat interval during performance was 693 (15) ms. Using as reference R-peak the one immediately preceding the wrong keystroke (R_0_), we assessed changes in the heart rate at preceding (R_−3_, R_−2_, R_−1_) and following (R_1_, R_2_) R-peak positions. A similar analysis was performed for R-peaks surrounding correct keystrokes (Figure 1C). Before and after correct events, there was a general linear increase (slowing) of the RR both before and following the keystroke (p < 0.05, FDR-corrected, PS_dep_ in range 0.76 – 0.88). For pitch errors, we found that the interbeat interval immediately preceding errors was larger than the previous RR-interval (RR_0_ = 0.691 [0.008] ms, RR-1 = 0.689 [0.009] ms, p < 0.05, FDR-corrected, PS_dep_ = 0.76) but not significantly different from the post-error interbeat intervals. A comparison between the RR-interval of error and correct events demonstrated larger values following correct keystrokes at position +2 (p < 0.05, FDR-corrected, PS_dep_ in range 0.92). Additionally, we assessed the relative change between consecutive RR-intervals (RR-interval at n position relative to RR-interval at the preceding n-1 position), to test for error-minus-correct differences in interbeat changes independently of the general linear increase of RR-intervals found for correct trials (Figure 1D). This representation demonstrated a relatively small RR-interval change at −2 and −1 RR positions in error trials, as compared to the larger changes found in correct trials (p < 0.05, FDR-corrected, PS_dep_ in range 0.64 – 0.82). Immediately before error keystrokes, however, this pattern was altered as there was a sudden increase in the RR-interval relative to the earlier RR-intervals (significant change increase in error trials at position 0, just before the error keystroke: p < 0.05, FDR-corrected, PS_dep_ in range 0.84). Following correct and error key presses, there was a reduction in the RR-interval change relative to the change found at position 0, which can be observed as a negative change in Figure 1F (p < 0.05, FDR-corrected, PS_dep_ in range 0.76 – 0.85).

Thus, we found that pitch errors were associated with a generally faster HR, yet were preceded by a slowing down of the RR-interval immediately preceding the key press. Pitch errors were then followed by a larger heart rate (HR) acceleration relative to correct keystrokes.

MVPA performed using trial-based patterns containing the three RR-intervals preceding the keystroke (error, correct) demonstrated a significant classification accuracy on the single-participant level in 13/17 participants (p < 0.05 in those individual subjects) and a significant group-level population accuracy of 57% (p = 0.004). A similar analysis including additionally the RR-interval following the error and correct key presses demonstrated a drop in the prevalence of the effect, as the classification accuracy on the single-participant level was significant in only in six pianists (p < 0.05).

Thus, information about an upcoming error or correct key press was more consistently decoded from the HR patterns preceding the keystroke.

### Effects of the cardiac cycle on behavioral performance

The same percentage of keystrokes fell into bin1 and bin2, 45 (1) % and 41(2) %, respectively, and these percentages were not significantly different (p = 0.212, similar rates for error and correct events), demonstrating that key presses were not aligned with a specific phase of the cardiac cycle (see also Figure S1).

Pitch errors during the diastolic phase (bin2) as compared to those in the systolic phase (bin1) led to a significantly larger post-error slowing and reduced keystroke velocity at +2K (p < 0.05, FDR-corrected; Post-error slowing, mean IOI at +2K: 137 (6) ms in bin1 and 144 (9) ms in bin2, PS_dep_ = 0.69; Keystroke velocity: 66 (1) in bin1 and 65(1) in bin 2, PS_dep_ = 0.71). These effects were not found in the immediate keystroke following pitch errors (+K) or at +3K (p > 0.05). Further, when assessing correct note events, we found no significant dissociation between the performance data in bin1 and bin2. These findings support that the effect of the phase of the cardiac cycle on performance was limited to timing and loudness properties of error processing in the subsequent keystroke at position +2K.

Changes in the RR-interval following errors did not differ when separately assessing error key presses in the cardiac systole or diastole (p > 0.05).

### Keystroke-locked ERP Analysis

The difference between the ERP waveforms for error and correct trials was statistically assessed with cluster-based permutation tests within [−100, 300] ms. We found a significant negative cluster preceding and a positive cluster following pitch errors (p = 0.0159 and 0.0199, respectively, p < 0.025, two-sided test). The negative cluster was due to a less positive pre-keystroke waveform in error trials than in correct trials within [−70, −30] ms (Figure 3A) with a frontal topography (Figure 3B; see also Figure S2, which includes the full baseline period in the visualization). The positive cluster had a frontocentral topography and was related to a positive post-keystroke deflection within [80, 300] ms in error trials (Figure 3C), which was not observed in correct trials (Figure 3D).

**Figure 3.**
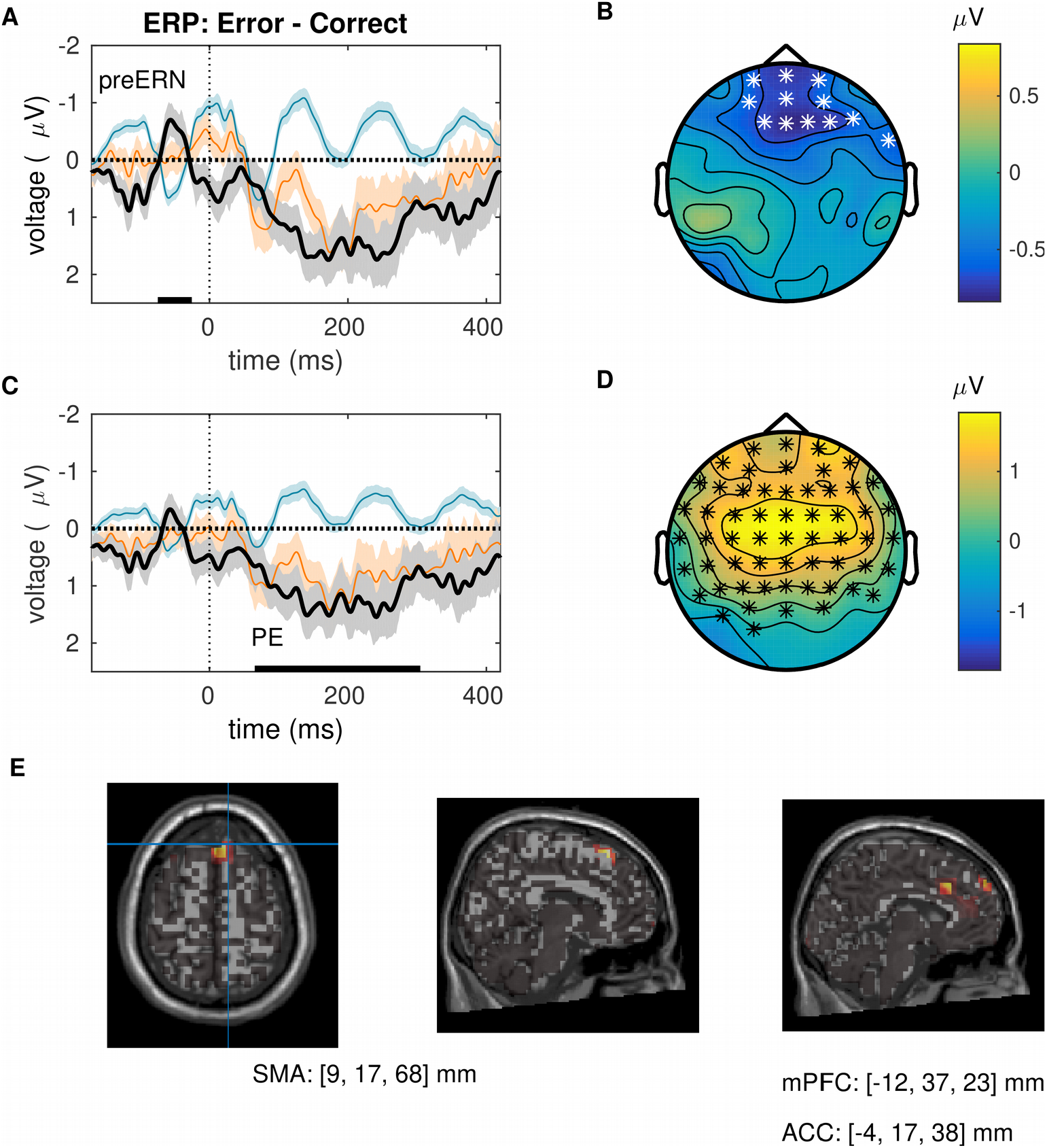
Neural signatures of error detection locked to keystrokes. **(A)**. Note-onset event-related potentials (ERP) depicted at frontal electrodes for pitch errors (orange line), correct notes (blue line) and for the difference (errors minus correct notes, black line). The keystroke at time 0ms is denoted by the dashed line. Shaded areas show ± 1 s.e.m. around the mean. The electrodes selected for this representation belonged to the significant negative cluster depicted in panel (B). Note the negative deflection preceding error commission (pre-error-related negativity, preERN). Black bars at the bottom indicate time windows of significant differences in this cluster (p = 0.0159, cluster-based permutation test). **(B)** Scalp topography for the preERN component, corresponding to the significant negative cluster obtained within −70 to −30 ms prior to keystrokes (p = 0.0159, cluster permutation test, p < 0.025, two-sided test) for the error-minus-correct note difference comparison. The white stars denote the electrodes belonging to the significant cluster. **(C)** ERP waveforms locked to keystrokes as in (A), but here averaged across significant electrodes that were part of the significant positive cluster represented in (D). The black bar at the bottom denotes the significant slower positive modulation following errors relative to correct key presses (error positivity, PE). **(D)** Scalp topography for the PE component, associated with the significant positive cluster obtained within 80 to 300ms following erroneous relative to correct keystrokes (p = 0.0199, p < 0.025). The black stars denote the electrodes belonging to the significant cluster. **(E)** Significant neural sources of the preERN (similar results for the PE: minimum norm estimate, regularization parameter *λ* = 0.1). MNI coordinates for sources at SMA, BA 6 (left panel), ACC and BA 9 in the medial prefrontal cortex (middle and right panels) are provided below.

Figure 3E displays the significant source localization outputs using minimum-norm estimates. In the time window of the preERN, there was a significantly smaller activation (p < 0.025, FWE-corrected) in a set of regions of the pMFC, such as the SMA (Brodmann area, BA 6, peak MNI coordinate at [X,Y,Z] = [9, 17, 68] mm), the ACC (BA 32 at [−4, 17, 38] mm), but also in the mPFC (BA 9 at [−12, 37, 23] mm). In the PE time window, significant differences in minimum-norm estimates were found in the ACC (BA 24 at [−7, 29, 18] mm) and the mPFC (BA 9 at [−10, 50, 48] mm; p < 0.025, FWE-corrected).

To exclude the potential confounding factor of different latencies in error trials driving the modulation of the preERN, as a control analysis we contrasted correct trials to a subset of error trials with matched latencies to the correct trials (see behavioral results: matched IOI with respect to the preceding keystroke). A cluster-based permutation test demonstrated a significant negative ERP deflection at −180 ms and a significant PE following errors around 150-350ms (Figure S3). A second frontocentral negative cluster was found, corresponding to the preERN, but showed a trend toward significance (p = 0.035, trend at the FWE-corrected level of 0.025). Thus, when differences in latencies preceding the error and correct keystrokes were controlled for, early error detection was associated with a negative deflection already 180 ms before the wrong keystroke and the preERN (trend) around [−80,−50] ms. As suggested by the main analysis, the PE was elicited following errors..

### ERP analysis locked to keystrokes in different phases of the cardiac cycle

We evaluated the keystroke-locked error-minus-correct difference ERP waveform separately for trials corresponding to bin1, 2 (systole, diastole). This analysis is similar to the one performed in the keystroked-locked ERP section (and Figure 3), but with a split of epochs according to the temporal interval (systole, diastole) in which the key following the R-peak was struck. A cluster-based permutation test performed within [−100, 300] ms revealed a significant negative difference between the error-minus-correct ERP waveform in systolic and diastolic phases of the cardiac cycle (p < 0.025, Figure 4). The effect reflected a more positive error-minus-correct ERP modulation for systole trials around 40-100 ms in a cluster of left centro-posterior electrodes. Based on the latency and topography, this effect seems to be dissociated from the error-related ERP components preERN and PE (Figure 3).

**Figure 4.**
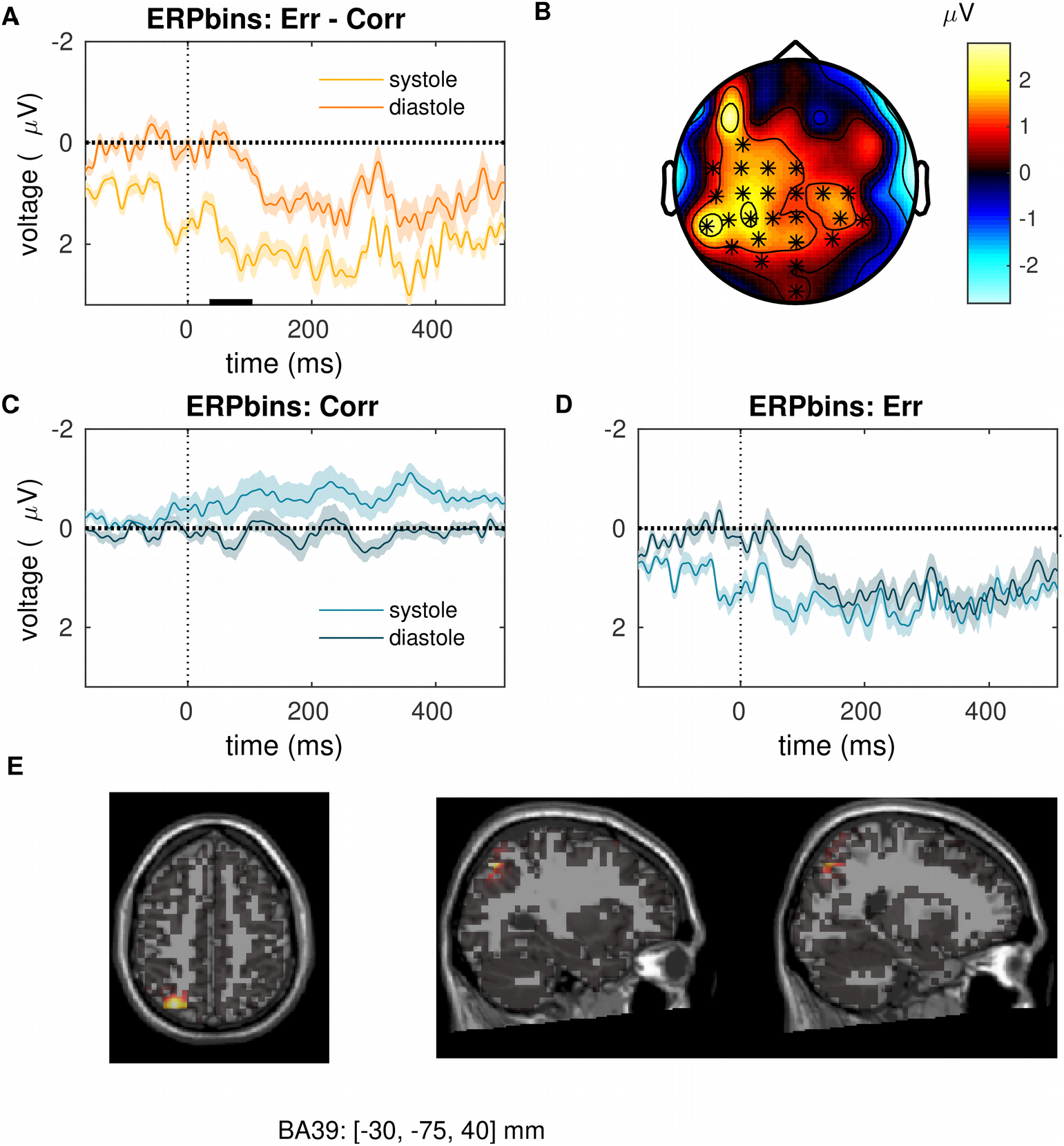
Influence of the cardiac cycle on the neural signatures of error detection locked to keystrokes. **(A)** Difference (error minus correct) event-related potentials (ERP) locked to keystrokes at time 0 ms (denoted by the vertical dashed line). The error-minus-correct difference ERP is plotted separately for bin1 and bin2 trials, in which the keystroke coincided with the systolic (light orange) or diastolic (dark orange) period of the cardiac cycle, respectively. The black bar at the bottom indicates the window of significant difference between the ERPs in the cardiac systolic and diastole periods (p = 0.021, two-sided cluster-based permutation test). The waveforms are averaged across electrodes pertaining to the significant cluster shown in panel (B). **(B)** Scalp topography for the significant cluster corresponding with the effect of the period of the cardiac cycle on the error-minus-correct ERP waveforms locked to keystrokes. Keystrokes coinciding with the systolic relative to the diastolic period elicited a larger positive deflection in the error-minus-correct ERP waveforms (p = 0.021, two-sided cluster-based permutation test). The black stars denote the electrodes belonging to the significant cluster. **(C)** ERP waveforms locked to correct keystrokes at systolic (light blue) and diastolic (dark blue) phases of the cardiac cycle. No significant difference was found between these two waveforms (p = 0.192). **(D)** Same as (C) but for pitch errors. No significant difference was found either (p = 0.281). **(E)** Significant neural source (minimum norm estimate, regularization parameter l = 0.1) for the difference between the error-minus-correct ERP waveform in the cardiac systole and diastole. Standard MNI coordinates of the anatomical location in the left inferior parietal cortex (Brodmann area 39) is shown below.

Of note was also that the time interval preceding the error events in bin1 and bin2 was not significantly different (the IOI relative to the previous keystroke was 147 [9] ms and 148 [7] ms in bins 1 and 2, respectively: p > 0.05). Thus, errors in the systolic or diastolic cardiac phase had similar temporal properties with respect to the previous keystroke, and they led to differences in the subsequent behavioral alterations later at +2K, that is, around 280 (10) ms following key presses (See *Effects of the cardiac cycle on behavioral performance*). Accordingly, the significant ERP effect within 40-100ms cannot be accounted for by a cardiac phase effect on the temporal properties of key presses within that temporal interval.

Source localization analysis demonstrated a significant difference between the systolic and diastolic modulation of the error-minus-correct ERP, with significantly increased cortical activation in the left inferior parietal cortex for systole trials (left BA 39 at [−30, −75, 40] mm).

### Multivariate Pattern Classification Analysis of ERP waveforms

Next, we used MVPA to assess whether the difference between the two phases of the cardiac cycle could be decoded from the patterns of single-trial ERP waveforms locked to pitch errors. For this analysis single epochs were locked to pitch error keystrokes falling either within the systolic (bin1) or diastolic (bin2) phase of the cardiac cycle. MVPA was evaluated at each time bin using as features the ERP amplitude values in the set of 64 channels.

Significant above-chance classification accuracy was found on the participant level preceding and following the pitch error in 15/17 participants (p < 0.05, FDR-corrected in all cases after control of FDR. Figure 5). On the group level, MVPA revealed a significant above-chance classification accuracy within [−86, −66] ms, [53, 73] ms and later within [173, 213] ms (p < 0.05, FDR-corrected, PS_dep_ = 0.970). Note that the second window converges with the cardiac cycle-based ERP effect reported in the previous section (*ERP analysis locked to keystrokes in different phases of the cardiac cycle*). These results indicate that the patterns of EEG activity across the cluster-related electrodes encoded class-related information, i.e. distinguishing between erroneous keystrokes coinciding with the systolic or diastolic phase of the cardiac cycle. A similar analysis performed with correct trials rendered no significant decoding accuracy at the group level (p > 0.05, mean accuracy 50%), but a significant single-participant decoding accuracy in 12 /17 subjects around −100 ms prior to the keystroke and after 200 ms (p < 0.05, FDR-corrected in all cases after control of FDR). Thus, the multivariate patterns of ERP waveforms were discriminative of the phase of the cardiac cycle more consistently for error trials. Moreover, primarily in error trials was there a result of significant decoding accuracy after 0 ms and before 100 ms, a time window that encompasses the early error-minus-correct ERP modulation for systole versus diastole. Errors in the systolic or diastolic phase had similar temporal properties with respect to the previous keystroke and at the following event at +K, yet they differed at +2K. Accordingly, the MVPA results obtained in the time windows preceding ([−86, −66] ms) and following ([53, 73] ms) the keystroke cannot be accounted for by timing differences in the two classes of error events.

**Figure 5.**
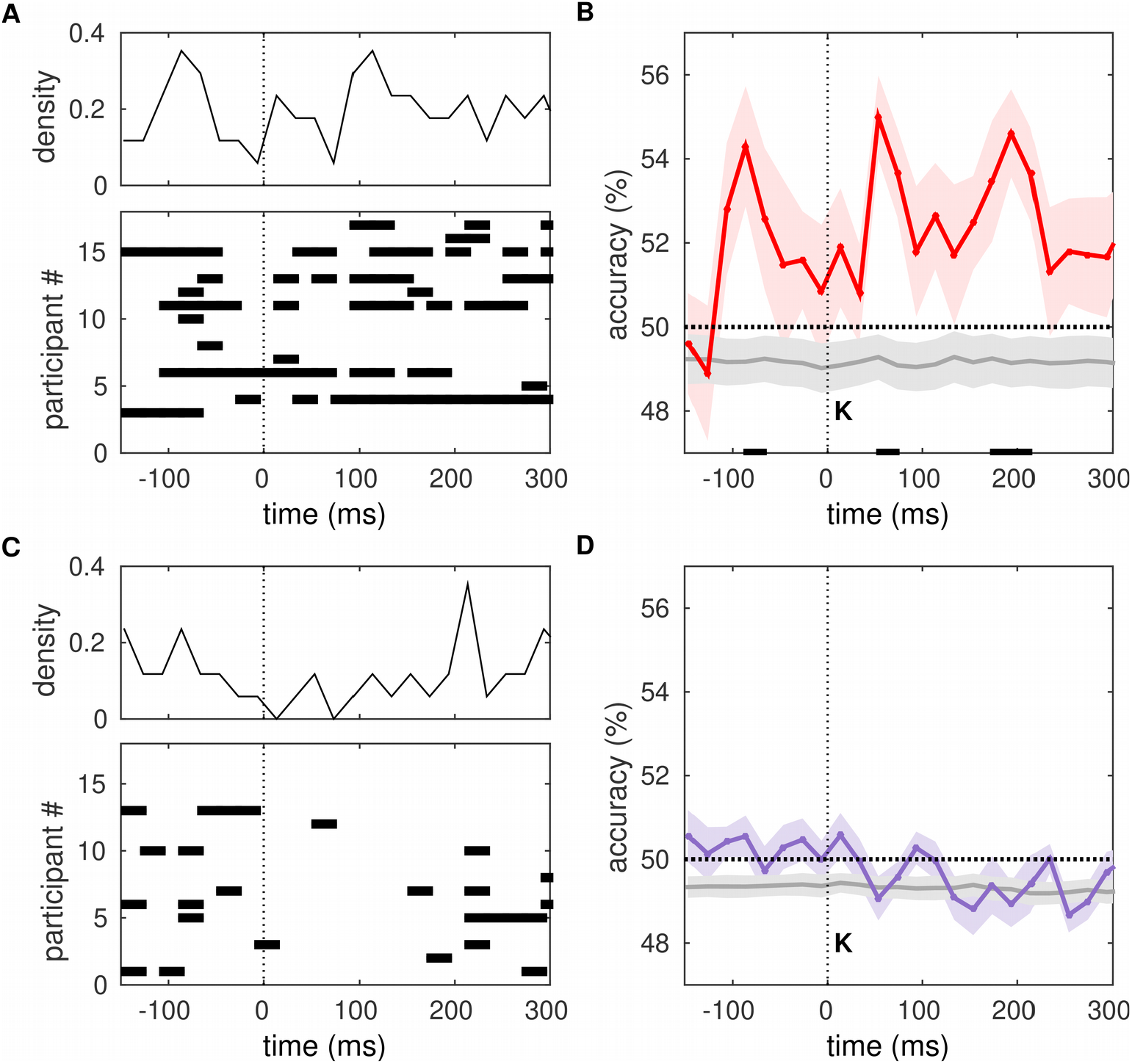
Multivariate pattern classification analysis (MVPA). MVPA for two classes of events: EEG trials locked to keystrokes in the cardiac systole and diastole. **(A)** Bottom: Time windows of significant above-chance decoding accuracy in error trials for single-subject MVPA analysis are denoted by horizontal bars (permutation tests, p < 0.05, FDR-corrected after control of FDR at level q = 0.05). Top: Density function representing the normalized total number of subjects showing significant classification accuracy in a time bin. **(B)** Population decoding accuracy for error trials plotted as the mean across participants (red line) and s.e.m. (red shade). Gray solid line indicates the chance level (s.e.m. as shaded gray area) on the group level estimated with permutation tests. Significant windows of above-chance decoding accuracy are denoted by the bottom vertical lines (p < 0.05, FDR-corrected, PS_dep_ = 0.970). The keystroke at time 0ms is indicated by the vertical dashed line. **(C)** Same as (A) but in correct trials. Significant above-chance decoding accuracy on the single-subject level is denoted by the horizontal black likes (permutation tests, p < 0.05, FDR-corrected). **(D)** Population decoding accuracy for correct trials plotted as the mean across participants (purple line) and s.e.m. (purple shade). There was no significant decoding accuracy above chance level in error trials (p > 0.05). The chance level (gray line; s.e.m shaded gray area) was estimated using permutation tests.

### Additional Control Analyses

To control for the possibility that differences in the ERP waveforms in each cardiac phase bin were accounted for by differences in the concurrent ECG modulations, we tested for statistical differences between the ECG signal locked to keystroke events in bin1 and bin2 (Figure 1D-E). In correct trials, the keystroke-locked ECG differed significantly between bin1 and bin2 within [−53, −8] ms preceding the key press (p < 0.05, FDR-corrected; PS_dep_ = 0.875). In error trials, the shifted ECG waveforms also differed significantly between bin1 and bin2 trials in the pre-keystroke interval within [−74, −18] ms (p < 0.05, FDR-corrected; PS_dep_ = 0.818). Thus, differences between the ECG signals in bin1 and bin2 conditions cannot account for the post-keystroke effects found with the cluster-based permutation test around 40-100ms. Neither can these differences be the source of the post-keystroke MVPA results at [53, 73] ms. However the MVPA effects found preceding the keystroke could be influenced by the ECG differences and should be taken with care.

In sum, the different behavioral and ECG control analyses support that the results from the ERP and MVPA analysis for bin1 and bin2 events found in the post-keystroke interval within 40-100 ms (ERP) and 53-73ms (MVPA) primarily reflect a cardiac-cycle-related influence on early error-monitoring processes.

## Discussion

This study sought to determine the role of cardiac afferent information in modulating early stages of error-monitoring during highly trained performance. In addition, we aimed to replicate the emergence of an anticipatory error detection ERP, the preERN, and a later PE during error commission in this setting, as reported in previous studies (Maidhof et al., 2009; Ruiz et al., 2009). Our results revealed that fluctuations in the cardiac cycle influence neural responses related to an early stage of error processing during action monitoring. Precisely, we found that the error-minus-correct ERP waveform had a more pronounced positive deflection within 40-100 ms for keystrokes that coincide with the systolic period of the cardiac cycle when compared with the cardiac diastole. This effect was dissociated in both time and topography from the specific error detection and evaluation components, the preERN and PE, respectively (Maidhof et al., 2009; Ruiz et al., 2009). Moreover, we revealed that the information about the systolic and diastolic phase of the cardiac cycle could be decoded from patterns of error-related neural activity on the single-trial and subject levels. In addition, we found that pitch errors in the systolic phase led to a smaller post-error slowing (PES) and less pronounced keystroke velocity reduction at position +2K. These findings support that during systole, when baroreceptor afferent activity is maximal, the neural responses to error processing are more pronounced, whereas the behavioral alterations following errors are attenuated.

The finding of an enhanced modulation of error-related neural responses during the cardiac systole is comparable to the previously reported effect of this cardiac phase on amplifying fear, pain and threat-related processing (Gray et al., 2012; Garfinkel et al., 2014). Several studies have investigated the impact of the dynamics of baroreceptor afferent activity on neural and psychological processes (Motyka et al., 2007; Gray et al., 2011; Garfinkel et al., 2014; Fiacconi et al., 2016; Azevedo et al., 2017). Baroreceptors are mechanoreceptors located in the heart and major blood vessels, which encode fluctuations in the cardiac cycle, such as the timing and amplitude of each heartbeat (Fadel et al., 2003). At the cortical level, visceral afferent representations – including those derived from baroreceptor activity – converge in the insular cortex, where they are integrated with pain and temperature sensations to guide behavior (Bennarroch, 1997; Critchley and Garfinkel, 2018). Previous studies have shown that during the cardiac systolic phase, there is a more pronounced processing of painful and fearful stimuli, threat appraisal, and expression of racial stereotypes (Gray et al., 2012; Garfinkel et al., 2014; Azevedo et al., 2017). By contrast, during the cardiac diastole, in which baroreceptor activity is minimal, detection of somatosensory stimuli is facilitated (Motyka et al., 2007; Al et al. 2018), and pre-motor responses to sensory stimulation are faster (Edwards et al., 2007). Thus, baroreceptor afferent activity modulates brain-body interactions during processing of sensory and painful stimuli, with a facilitatory role of maximum baroreceptor activity on threat-related processing.

In light of these findings, our results support that errors are salient events whose processing relies not only on proprioceptive and exteroceptive information, as shown previously (Ruiz et al., 2009; Orr and Carrasco, 2010; Boldt and Yeung, 2015; Ullsperger et al 2014), but also on implicit interoceptive cues, such as enhanced cardiovascular afferent signals. Ullsperger and colleagues (2010) have proposed that, following initial error detection by the pMFC, the conscious awareness of an error – as signalled by the PE component – engages the saliency network in addition to the executive network. Interestingly, the anterior insula (AI) and ACC are part of both networks (Seeley et al., 2007; Menon and Uddin, 2010). Moreover, the AI and ventral ACC are key centres for processing visceral information (Park et al., 2014; Barrett and Simons, 2015; Hassanpour et al., 2018). The AI has been linked to the generation of the PE and modulation of error awareness (Dhar et al., 2011). Additional correlational evidence coming from reaction-time tasks has shown that the PE is modulated by explicit interoception (heartbeat counting task, Sueyoshi et al., 2014). These studies thus favour the interpretation that interoceptive processing interacts with error monitoring mainly during the later stages, contributing to the conscious awareness of an error.

Crucially, we found that the error-minus-correct ERP modulation by cardiovascular interoceptive signals occurred within 40-100ms, and thus preceded the PE component, which peaked around 150-250ms. This novel finding expands previous results (Dhar et al., 2011; Sueyoshi et al., 2014), by suggesting an earlier window in which visceral interoceptive information can modulate error processing, before conscious experience of the sensory consequences of an action. This finding converges with the recent data on an earlier involvement of the AI – and potentially interoceptive processing – in error detection (Bastin et al., 2017). Thus early changes in cardiovascular information during error-monitoring processes could shape subsequent conscious evaluation of the error, thereby modifying ongoing performance. This interpretation is in agreement with a predictive account of interoception, whereby predictions about upcoming visceral sensations can drive behavioral and physiological responses to maintain homeostasis (Barrett and Simons, 2015; Seth and Friston, 2016; Tsakiris and De Preester, 2019). Predictive interoceptive processes are modulated by activity in the anterior insula (Barrett and Simons, 2015), and can shape ongoing behavior and (neuro) physiological states (Hassanpour et al., 2018), including error-monitoring processes (Bastin et al., 2018).

More generally, the predictive account of interoception can be framed within the context of predictive coding and its formulation in terms of Bayesian active inference principles (Seth, 2013; Barret & Simmons, 2015). According to the influential active inference account, the brain continually generates predictions about sensory inputs based on prior experience (Friston, 2010; Friston et al., 2018; Tsakiris and De Preester, 2019). These predictions are based on probabilistic models that map (hidden) causes of sensory inputs and observed data (Friston, 2010; Clark et al., 2013). Active inference and predictive coding models are defined in hierarchical brain networks in which top-down signals carry predictions and bottom-up signals pass on prediction errors (the mismatch between expected and actual outcomes). Long-term experience, as in the case of skilled music performance or vocal production, can shape these predictions (Shum et al., 2011; Koelsch et al., 2019), thereby modulating neuronal activity in anticipation of the unwanted sensory consequences of ongoing movements associated with errors (Ruiz et al., 2009; Maidhof et al., 2009, 2013; Shum et al., 2011). Anticipatory changes in neuronal activity would aim to minimize the error between the predicted and the actual sensation, which would trigger anticipatory corrective adjustments (Ruiz et al., 2011; Palmer et al., 2012; Shum et al., 2011; Behroozmand and Sangtian, 2018), as reflected here in the reduced loudness and slower tempo at the error and preceding keystrokes.

The effect of the cardiac cycle on the error-minus-correct ERP was linked to activity in the left inferior parietal lobe (IPL), a region implicated in human cardiac autonomic regulation and visceral processing (Lutz et al., 2009; Park et al., 2014). By contrast, source localization of the preERN and PE components revealed regions of the pMFC, such as the SMA, ACC and BA 9, which is consistent with the well-documented involvement of the pMFC in general error-monitoring (Dhar et al., 2011; Ullsperger et al., 2014). The IPL, along with the ventral ACC, AI and somatosensory areas, is part of the visceroneural network, which receives visceral information about bodily states (Lutz et al., 2009; Kleckner et al., 2017). Activity in this network might trigger changes in heart rate, such as HR deceleration following perceptual misses or response errors (Park et al., 2014; Bastin et al., 2017). Here, we found a HR acceleration following pitch errors relative to correct keystrokes, which seems to stand in contrast with the post-error HR deceleration reported in previous studies (Hajcak et al., 2003; Wessel et al., 2011, Bastin et al., 2017). Significantly, however, a slowing down of the interbeat interval was found in error trials immediately preceding the key press. Moreover, the pattern of preceding HR changes at the single-trial level contained information predictive of whether an upcoming key press was an error or a correct event. The HR deceleration and acceleration effects before and after error commission, respectively, were independent of the cardiac phase with which errors coincided. Behaviorally, however, we found that following errors in the cardiac systole there was a significantly reduced slowing of tempo and keystroke velocity at +2K post-error. These results, therefore, suggest that the modulation of IPL activity by cardiac interoceptive signals during error processing might be associated with a tuning of the gain on anticipatory HR changes driving subsequent behavioral alterations following pitch errors.

The early involvement of the IPL in our study also aligns well with its role in multisensory integration, including proprioceptive and interoceptive signals, in anticipation of errors during sensorimotor adaptation for speech and motor tasks (Ghilardi et al., 2000; Luauté, et al., 2009; Shum et al., 2011). The IPL is reciprocally connected with the primary somatosensory cortex (S1, Borich et al. 2015) and S1 in turn with the somatotopically similar primary motor cortex, M1, integrating motor commands with proprioceptive and tactile information even before motor initiation begins (Bouchard et al. 2013). This evidence supports our conclusion that during the cardiac systole, an IPL-driven mechanism may enhance neuronal processes engaged in error-monitoring. In the context of self-generated errors, but also for feedback-based errors (such as auditory feedback alterations for speech), a higher activation of the IPL could indicate a more active salience network that aids the detection of inconsistencies related to processes occurring prior to the perception of unwanted sensory consequences.

Overall, we propose that a larger anterior IPL-dependent neural response during cardiac systole might reflect enhanced processing of interoceptive cues contributing to error processing. The anticipatory changes in cardiovascular information (sudden HR increase in error trials before the key press) could provide a form of feedback in addition to the early error-detection signatures to influence ongoing performance, and adapt subsequent behavior. The cardiac systole may amplify processing of these early interoceptive cues, thus facilitating error detection, thereby diminishing the window of integration over which evidence for an error could be accumulated. Consequently, the slowing of responses following the error in the systolic phase is less pronounced. Note that in our study, PES effects extended for six successive keystrokes – or three keystrokes when assessing pitch errors of similar pre-error temporal properties than correct key presses – until timing performance progressively returned to baseline levels, converging with previous studies (Ruiz et al., 2009; Maidhof et al., 2009; Palmer et al., 2012). In this context, re-establishing accurate timing following error commission might be crucial for skilled performance (Goebl and Palmer, 2013), which here was facilitated during systole. Our interpretation is consistent with new evidence supporting that post-error adjustments can be considered as a manifestation of a general orienting reflex rather than goal-directed adaptation (Notebaert et al., 2009; Wessell et al., 2017). As such, facilitation of error processing during maximal baroreceptor activity might ameliorate orienting responses and post-error slowing.

In conclusion, this study offers a novel account of implicit cardiovascular interoceptive signals modulating neural and behavioral responses related with early error monitoring during skilled performance. It also provides the first evidence that prediction of upcoming errors triggers not only anticipatory error-detection ERP components, such as the preERN (Ruiz et al., 2009, Maidhof et al., 2009), but also anticipatory changes in interoceptive states. HR changes triggered by errors have been taken as evidence for an influence of error-related interoceptive signals on subsequent perceptual processing (Lukowska et al., 2018). Phasic activity in the locus coeruleus– norepinephrine system could drive these cardiovascular changes in response to errors (Ullsperger et al., 2010; Lukowska et al., 2018). Our study extends the findings of Lukowska and colleagues (2018) to the anticipatory period, when predictive error-detection components can shape anticipatory HR responses, thereby influencing subsequent performance. Thus, the data suggest that prediction of the upcoming consequences of our actions relies on the integration of exteroceptive and interoceptive information, a process likely shaped by long-term training (Schirmer-Mokwa et al., 2015; Kleber et al., 2013), and involving higher order somatosensory regions and multisensory integration regions in the IPL.

Some aspects of the present study should be carefully considered when interpreting the results. First, the fast performance rate (125 ms between consecutive notes) imposed on the pianists to elicit pitch errors – which is realistic in highly skilled music performance – can lead to overlapping ERP components of neighbor events (Woldorff 1993). This limitation was addressed by Herrojo Ruiz et al. (2009) by complementing the ERP analysis with a second method aimed at disentangling possible overlapping brain responses, the symbolic resonance analysis (Beim Graben and Kurths 2003). A possible improvement on the current paradigm that would allow for the disentanglement of ERPs to different types of events with similar latencies is a longer recording session. A larger sample of error events of different latencies would allow for a detailed match of latency between error and correct events, while allowing for larger epoch numbers and therefore higher signal-to-noise ratio in this analysis. A second effect of the fast performance rate was the emergence of a cyclic pattern of ongoing changes in the ERP responses, corresponding to the preparation, execution, and evaluation of individual key presses. This should be taken into account when interpreting the differences between baseline-corrected ERP waveforms, as the baseline period in all conditions was affected by the modulation of ongoing motor responses.

An additional limitation of our study is that EEG and source localization do not allow the investigation of insular activity during the anticipation and processing of errors, which would be crucial to understand the mechanisms by which anticipatory interoceptive states drive early error processes. These are likely mediated by insular activity. In addition, we did not use individual MRI scans to inform source modelling, but a standard template anatomy, as validated elsewhere (Haufe et al., 2016). Accordingly the source localisation results should be interpreted with care and need to be replicated in future studies. Follow-up studies using combined MRI and EEG or magnetoencephalography to assess cardiovascular influences on error processing across different stages of training will be central to further our understanding of the impact of interoception in expert performance.

## Acknowledgements

The study was supported by Goldsmiths University of London, the British Academy, through grant SG161006, and the German Research Foundation (DFG - Deutsche Forschungsgemeinschaft), through project HE 6013/1-2 to MHR. The authors would like to thank two anonimous reviewers for their excellent suggestions, which improved the manuscript considerably.

## Author contributions

M.H.R. and J.B. designed the experiment, G.B. and M.G. collected the data, M.H.R. analysed the data, M.H.R., G.B. wrote the manuscript, J.B. edited the manuscript.

## Supplementary Figures

**Figure S1.**
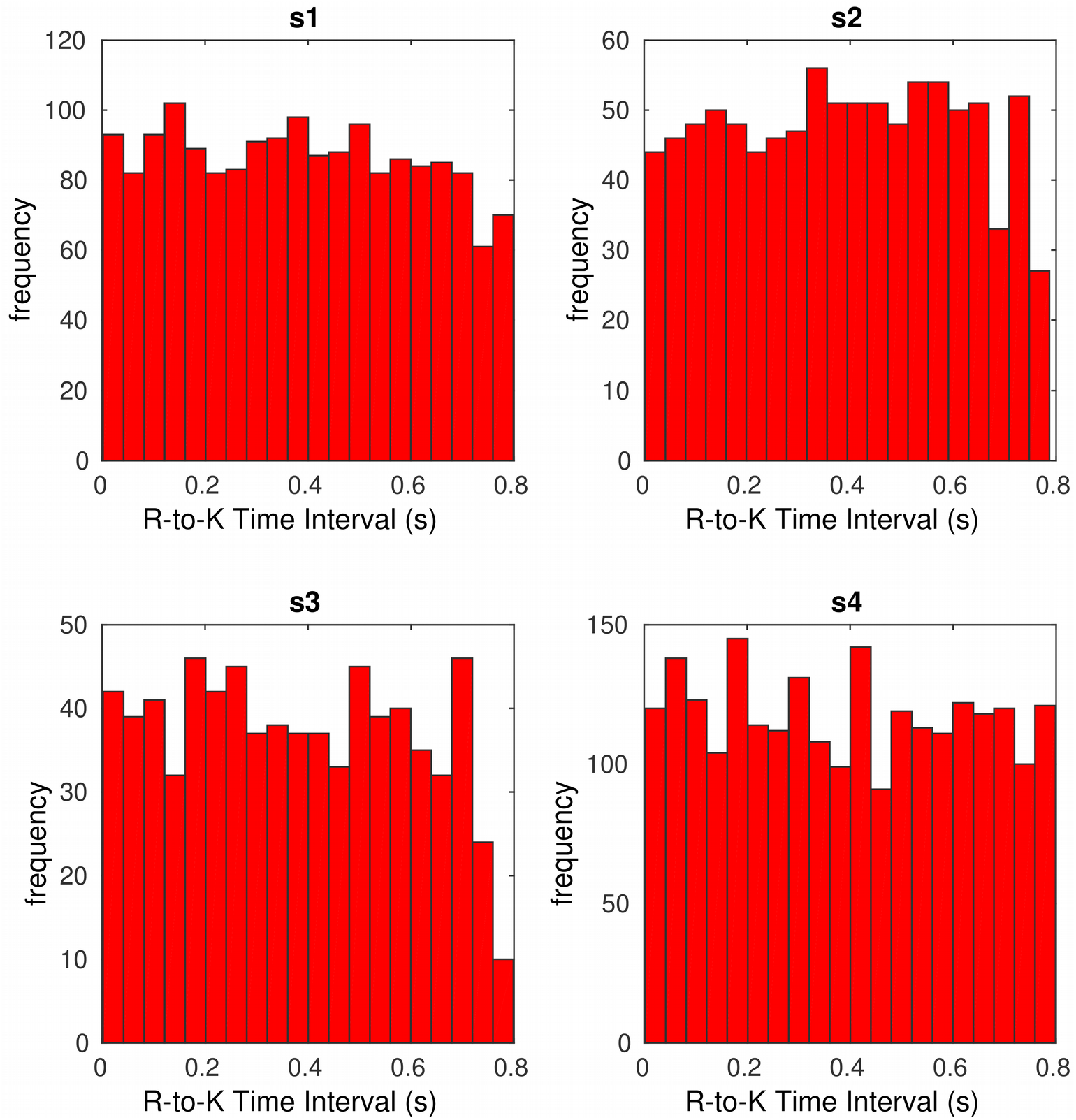

**Figure S2.**
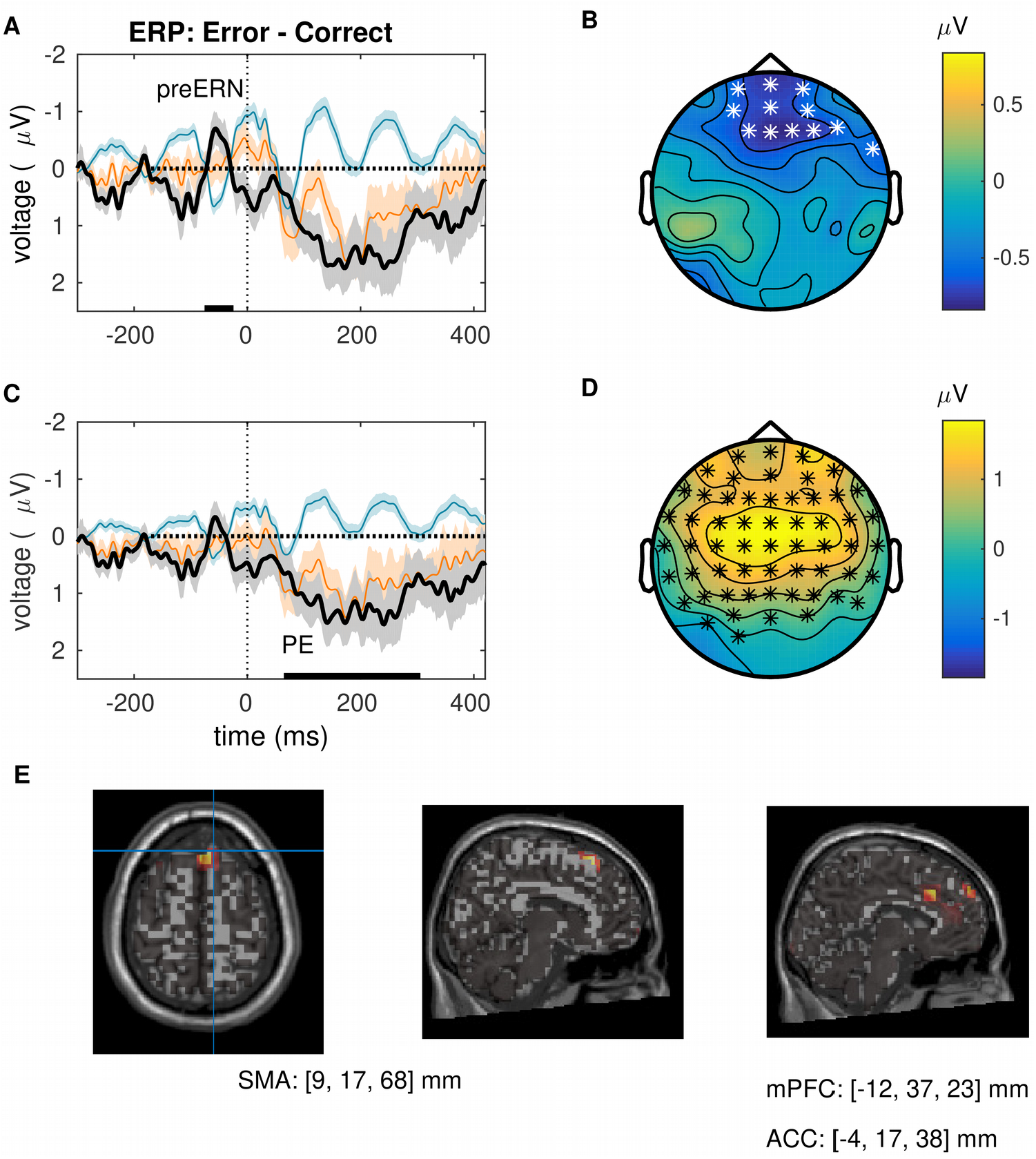

**Figure S3.**
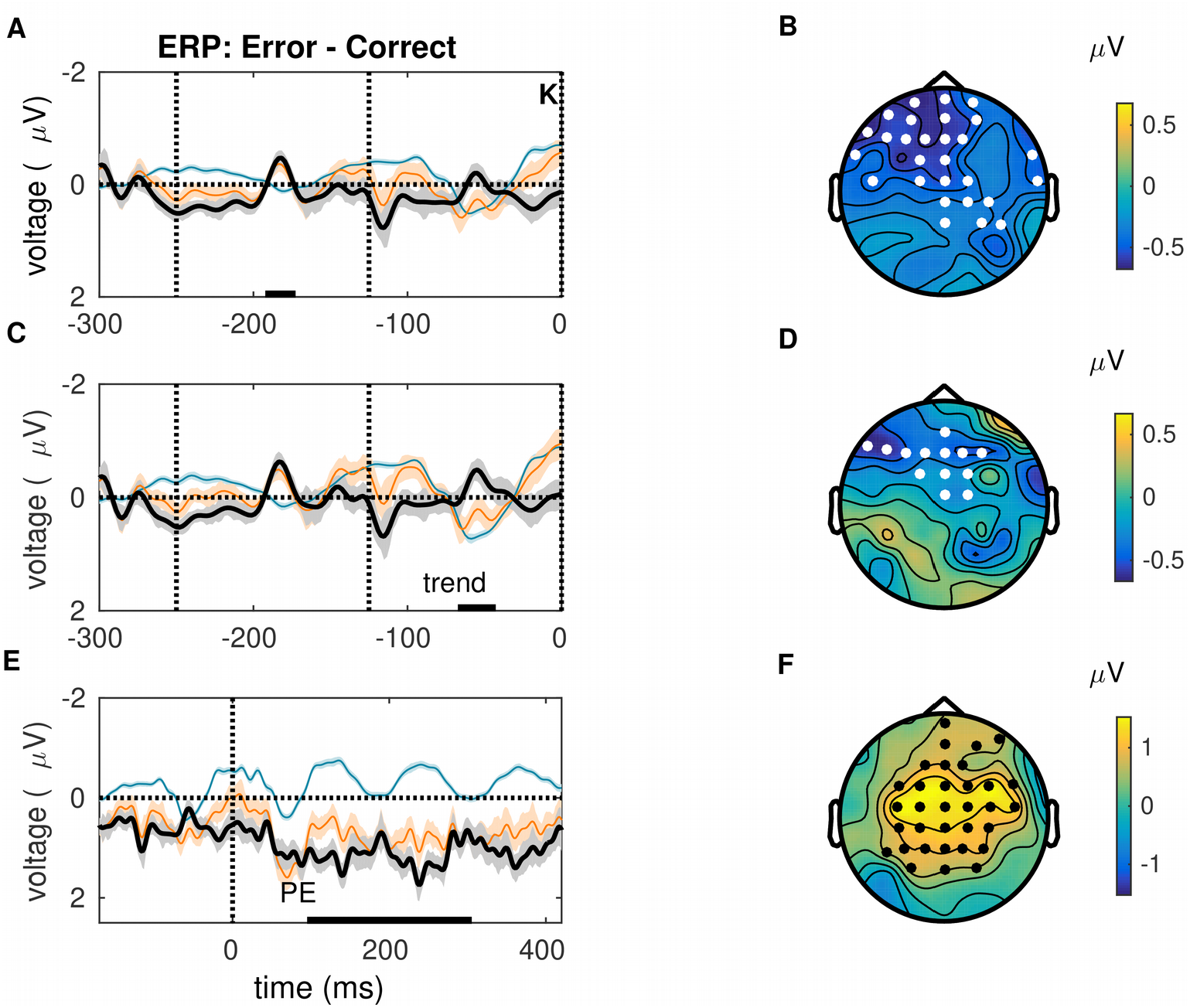

